# Age of seizure onset as a major determinant of memory changes following surgery for hippocampal sclerosis

**DOI:** 10.1101/800888

**Authors:** Eduardo Leal-Conceição, Marino Muxfeldt Bianchin, Wyllians Vendramini Borelli, Liss Januário de Oliveira, Mário Bernardes Wagner, André Palmini, Eliseu Paglioli Neto, Graciane Radaelli, Vinícius Spencer Escobar, Jaderson Costa da Costa, Mirna Wetters Portuguez

## Abstract

**Objective:** To study the relation of epileptological and surgical variables with post-operative memory performance, following surgery for refractory mesial temporal lobe epilepsy (MTLE) due to hippocampal sclerosis (HS).

**Methods:** Logical memory (LM) and visual memory (VM) scores for immediate and late recall of 201 patients operated for MTLE/HS from 1996 to 2016 were reviewed. All patients were evaluated prior to surgery and reevaluated up until 5 years after the procedure. Scores were standardized to a control group of 54 healthy individuals matched for age and education. Patients were divided in two groups according to the hemisphere affected and scores for immediate and late recall were compared. Reliable Change Index (RCI) with a 90% confidence interval was performed to verify individual memory changes for each late LM (lLM) and late VM (lVM) score. A multiple linear regression was performed with the RCI using late recall scores of lLM and lVM and clinical variables.

**Results:** 112 (56%) patients had left HS (lHS). The lHS group showed decreased immediate LM (iLM) scores before and after surgery (p<0.05), compared with rHS. The rHS group showed increased iLM scores post-operatively (p<0.05). RCI of the rHS group showed that 6 (7%) individuals had improved, 78 (87%) stabilized and 5 (6%) decresead in lLM scores, and for lVM 7 (8%) improved, 80 (89%) stabilized and 2 (3%) worsened (RCI> 1.645). RCI of the lHS group showed that 3 (3%) individuals had increased scores, 104 (93%) stabilized and 5 (4%) worsened for lLM, and for lVM 3 (3%) obtained improved scores, 103 (92%) stabilized and 6 (5%) decreased (RCI> 1.645). lHS and later age of onset of the first epileptic seizure were predictors of lLM loss (p<0.05).

**Conclusion:** Left MTLE/HS and later age of seizure onset were predictive factors for worsening of lLM. No RCI predictors of post-op lVM performance were identified. We found a decreased baseline functioning of LM in the lHS group and improvement of LM in some patients who had resection of the right MTL. Patients of the rHS group had a higher percentage of reliable post-op improvement for both VM and LM scores.

## INTRODUCTION

MTLE/HS is the most prevalent epileptic syndrome which is both refractory to medication and surgically-remediable^1,2^. Despite high rates of seizure control, resection of mesial temporal structures obviously impinge upon memory circuits, and therefore post-operative memory decline is a major concern.

Psychometric testing, combined with clinical interview, has the potential to identify the individual’s basal cognitive functioning, the hemispheric lateralization of cognitive dysfunction and the functional performance of the region to be removed^3^. Also, when evaluation is performed pre and post-operatively, the impact of the procedure on individual’s cognition can be assessed^3^. However, despite numerous studies, it is still unclear which patients are at higher risk of significant memory decline following resection. For instance, although there is enough evidence linking left lateralization of the disease and verbal logical memory, it is difficult to determine in an individual basis the risk of significant decline, even taking into consideration the presence of HS and preoperative functionality^4,5^. Thus, a degree of uncertainty often compromises decision on the best surgical strategy, and research is certainly needed on prognostic factors for decline^6^.

Predictive factors of post-op memory changes vary widely among studies and include surgical technique, age at surgery, preoperative cognitive status and seizure freedom. Selective resection of mesial structures in patients who already have significant memory abnormalities, being seizure free, and an early age at surgery appear as protective factors against significant declin^7–10^. However, these findings are not universal^7,11^ and such discrepancies prevent solid decision making in an individual basis. Important reasons for heterogeneous findings are that many studies include patients with a broad range of pathologies impinging upon different structures of the MTL and methodologies to identify post-operative changes in memory are distinct^12^. Here, we present the correlation of a large number of clinical, epileptological and surgical variables with reliable memory changes after 1 to 5 years following resection of a single, homogeneous pathology in the MTL. Restricting the analyses to patients with HS, the most prevalent subtype of MTLE, allowed us to provide a more reliable perspective of predictors for changes in memory performance.

## METHODS

### Subjects

**Data** was collected from medical records of 201 patients with ages between 16 and 60 years, evaluated and operated for MTLE/HS at the Epilepsy Surgery Program of Hospital São Lucas of PUCRS, between 1996 and 2016. MRI fidings suggestive of HS were independently confirmed by an experienced neuroradiologist and by the epileptological team. Individuals were then subdivided in two groups according the hippocampal sclerosis lateralization: left or right. : anterior temporal lobectomy (ATL) or selective amygdalohypocampectomy (SAH).

### Data acquirement

Sociodemographic, clinical and neuropsychological data was collected for analysis. The Engel scale^13^ was performed to evaluate the impact of surgery on the frequency of the seizures after surgery. All individuals underwent a comprehensive neuropsychological testing before and after the neurosurgical procedure within a 5-year interval, including the Wechsler Memory Scale – Revised^14^. We collected the memory scores of each individual in the logical memory and visual memory recall tests (LM and VM respectively). In the LM test, two stories are read to the patient and the patient must recall the reading immediately (iLM) and 30 minutes later (lLM). The VM test consists in the presentation, during 10 seconds, of 4 cards with geometric figures - one at a time. The patients need to draw what they remember of the images immediately (iVM) and 30 minutes after the presentation, corresponding to the late score (lVM). The survey and punctuation were performed according to instructions.

### Statistical Analysis

Both group and individual analysis were performed in this study. Parametric tests were used to compare sociodemographic and memory scores between groups (Figure 1). Individual analyses were performed according to the Reliable Change Index (RCI) a method developed by Jacobsen and Traux^15,16^ and later modified by other psychometrists^17–20^ (Figure 1). The RCI is the gold standard to evaluate cognitive alterations after any kind of intervention, because of its efficiency in withdraw the effects of practice, learning and measurement errors in the postoperative evaluation^15^. For the calculation, the control population from the WMS-R^14,21^ manual was used to normalize scores in standard deviation (SD), in comparison to the healthy population, and a confidence level of 90% was considered^17,19,20^. For RCI, only lLM and lLV data were used, considering the importance of temporal mesial structures for this type of function^22^. The results of the RCI’ are presented in percentage, classified as improvement (RCI>+1.645), stability (−1.645< RCI <1,645) or worsening (RCI <−1.645) of function. Multiple linear regressions were also performed to find predictors of memory changes. All statistical analysis were performed with the RStudio program (v1.0.136)^23^, and it was considered a p<0.05 statistically significant. All patients have the same underlying disease and this index will be used in order to reduce the possibilities of biases and measurement errors in relation to the memory change.

**Figure1:**
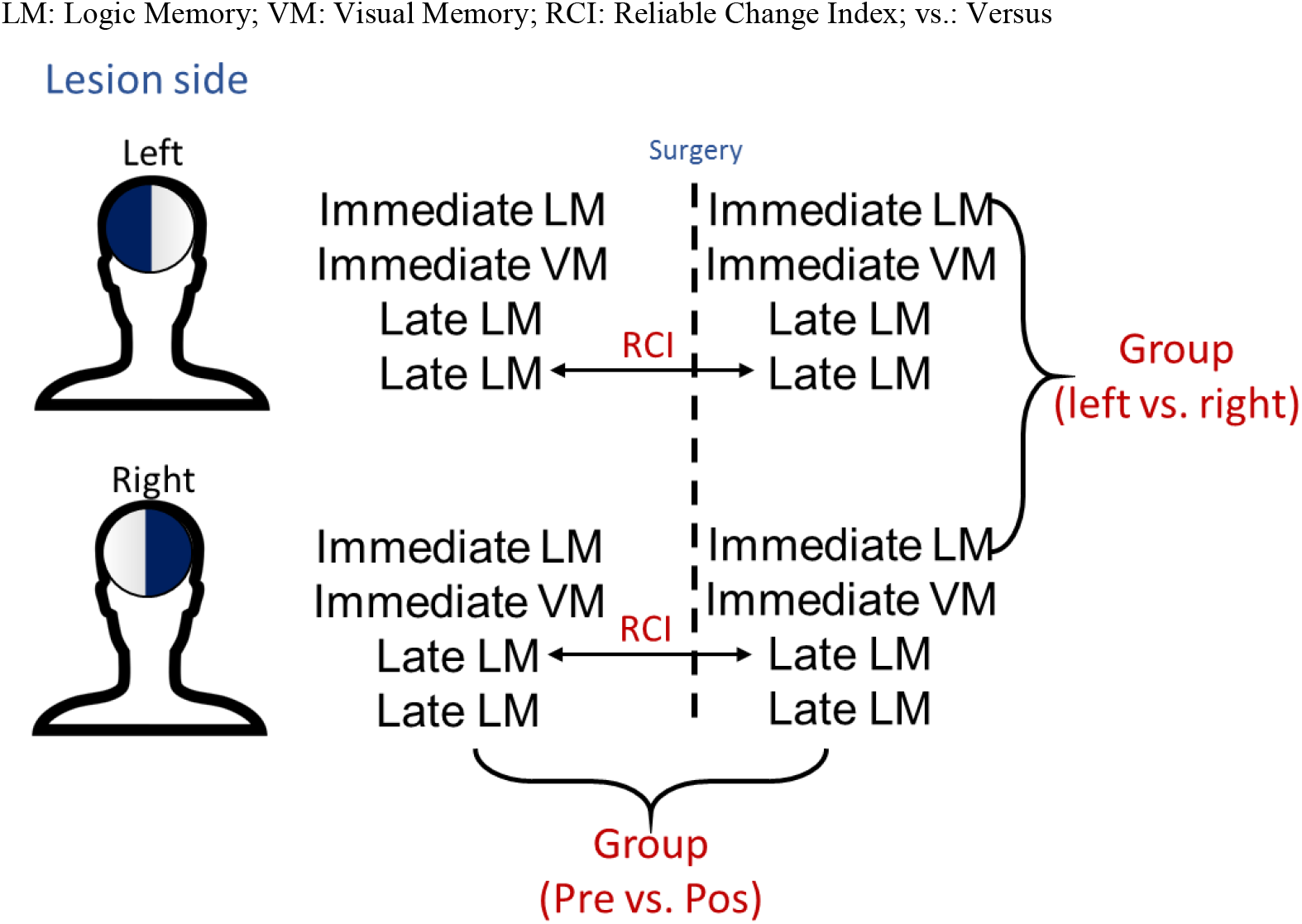
Model used to compare memory results.

## RESULTS

### Sample characteristics

Both lHS and rHS groups shared similar sociodemographic characteristics according to age, sex, education (Table 1). The lHS group showed lower baseline lLM scores when compared with the rHS group both in preoperative (13,06+−8,10 vs. 16,03+−7,9), p=0.0049) and postoperative (12,18+−7,65 vs. 18,45+−8,91, p<0.0001) settings (Table 2). However, when comparing the performance in the neuropsychological evaluation between the two groups, a statistically significant difference was found in the results of preoperative (p = 0.0049) and postoperative (p = 0.0001) lLM, demonstrating a lower performance of these functions in patients with lesions in the Left Hemisphere (LH). Regarding lVM, no significant differences were found (p> 0.05):

**Table 1:**
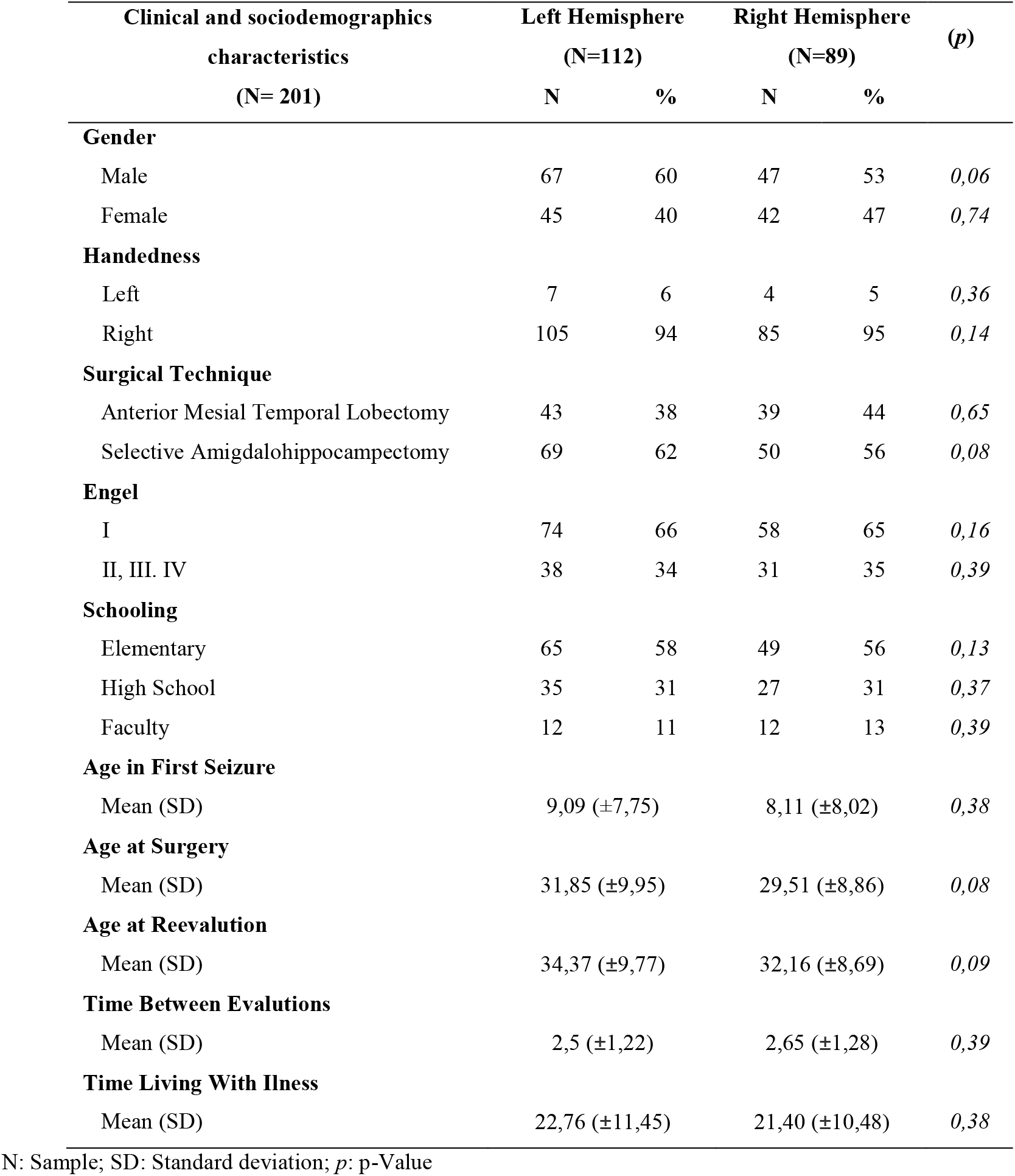
Sociodemographic, clinical and neuropsychological information.

**Table 2:**
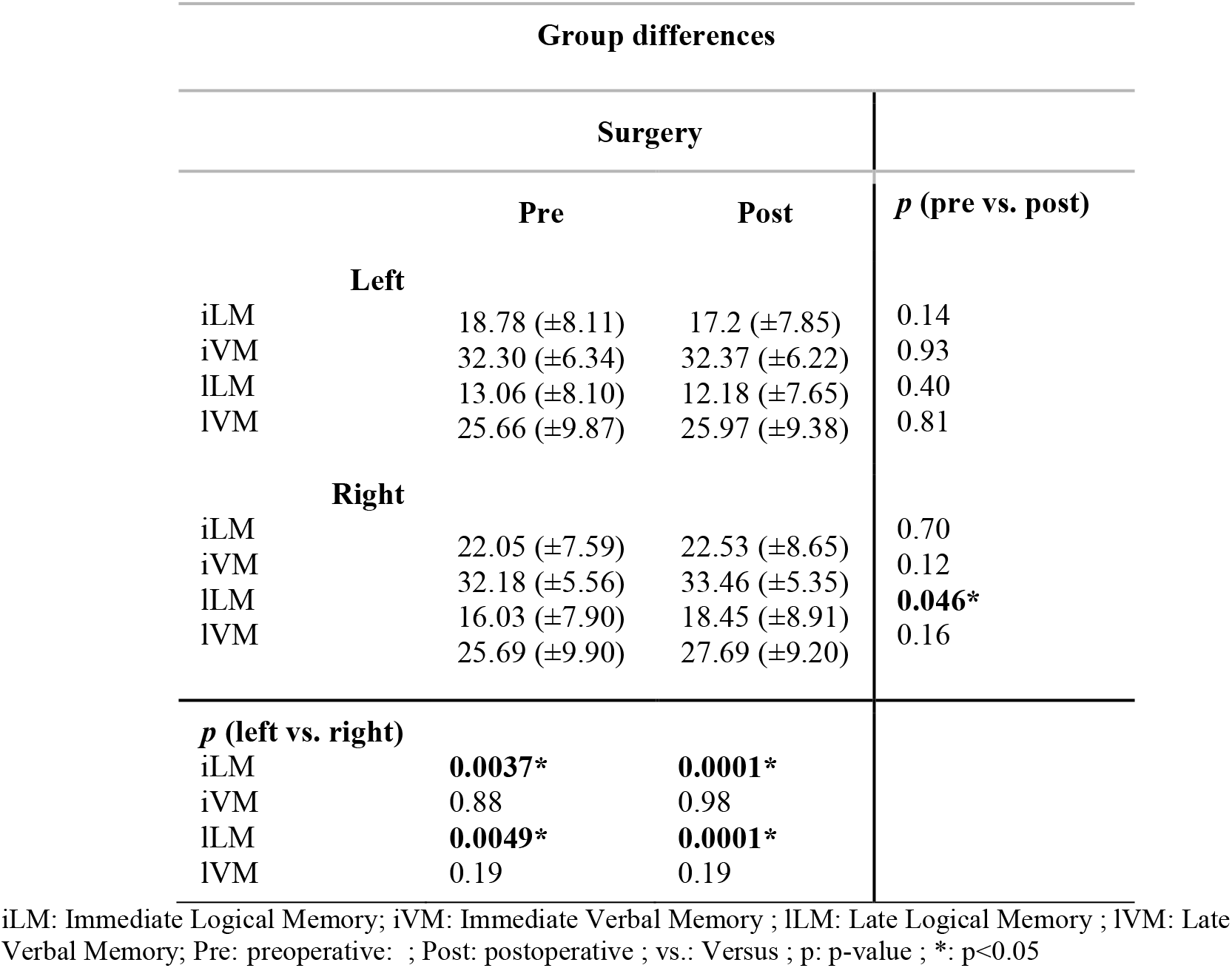
Neuropsychological results and comparison between group performances.

### Between and within group differences

Both groups shared similar type of surgery, Engel scores, level of education, time for the first seizure, time for reevaluation and time for living with the disease (p>0.05 for all measures) (Table 1). Subjects of the lHS group showed a mean time of 2.5 (± 1.22) years between baseline evaluation and retesting, while the rHS group showed a mean of 2.65 (± 1.28) years (p>0.05). When compared with the lHS group, the rHS group showed significantly higher scores in the iLM test in the baseline (22.05±7,59 vs. 18.78±8,11, p=0.0037) and after surgery (22.53±8,65 vs. 17.2±7,85, p=0.0001). The rHS group also showed increased lLM scores than the lHS subjects before (16.03±7,9 vs. 13.06±8,10, p=0.004) and after surgery (18.45±8,91 vs. 12.1±7,65, p=0.0001). (Table 2, Figure 1) However, the VM scores did not differ between groups either before or after the surgical procedure (p>0.05 for iVM and lVM measures). Regarding within group differences, the rHS group showed a borderline increase in lLM scores when comparing before and after surgery (16.03±7,9 vs. 18.45±8,91, p=0.046). All other LM and VM scores did not show a significant difference within group analysis comparing before and after surgery either for lHS or rHS group (p>0.05 in all other measures) (Table 2).

### Individual memory changes

Regarding the rHS group, 55 subjects showed decreased scores of lLM on baseline and 61 subjects on follow-up when compared with the normative data from the population. Besides, 37 individuals showed decreased lVM scores on baseline and 37 on follow-up. The RCI of the lLM scores of the rHS group showed that the majority of individuals were stable after surgery (104 subjects, 93%), while 3 (3%) subjects improved and 5 (4%) worsened. For the lVM scores of this group, the majority of individuals also were stable after surgery (103, 92%), but 3 improved and 6 worsened during this period (3% and 5% respectively) (Table 3).

**Table 3:** Results of individual memory changes.

| Individual differences (RCI) |  |  |  |
| --- | --- | --- | --- |
|  | Improved | Stabilized | Decreased |
| **Left** |  |  |  |
| ILM | 3 (3%) | 104 (93%) | 5 (4%) |
| IVM | 3 (3%) | 103 (92%) | 6 (5%) |
| **Right** |  |  |  |
| ILM | 6 (7%) | 78 (87%) | 5 (6%) |
| IVM | 7 (8%) | 80 (89%) | 2 (3%) |

ILM: Late Logical Memory ; IVM: Late Verbal Memory

In patients with rHS, compared with normative data from the control population, 55 patients had a deficit related to late LM on preoperative period and 61 on postoperative period. For late VM, 37 subjects presented deficit scores in the evaluation that preceded the surgery and 33 in the posterior testing.

When calculating RCI for this group of patients, with a confidence interval of 90%, 3 (3%) of the patients presented improvement, 104 (93%) stability and 5 (4%) worsened in relation to late logical-verbal memory. For late visual memory, 3 (3%) showed improvement, 103 (92%) stability and 6 (5%) worsening (Table 3).

In relation to the lHS group, 40 individuals showed lower scores of lLM in baseline and 30 individuals on follow-up when compared with the normative data from the population. Besides, 31 individuals showed decreased scores of lVM in the baseline and 21 individuals on follow-up also compared with normative data. The RCI of the lLM scores in the lHS group showed that the majority of individuals were stable after surgery (78 subjects, 87%), while 6 improved and 5 worsened (7% and 6% respectively). Regarding the lVM scores, again the majority showed stabilized scores (80 subjects, 89%), while 7 improved and 2 worsened (8% and 3% respectively) (Table 3).

In patients with lHS, comparing the scores of this group of patients with a healthy control population, 40 individuals had a poorer performance in relation to late LM in the previous and 30 in the posterior evaluation. Considering late VM, 31 showed deficits in the preoperative testing and 21 in the postoperative.

Using the method of RCI, it was evidenced that approximately 6 (7%) of the patients presented improvement, 78 (87%) stability and 5 (6%) worsened in lLM scores. In this same group, 7 (8%) showed improvement, 80 (89%) stability and 2 (3%) worseneding in relation to lVM (Table 3).

### Multiple Linear Regression for Memory

The linear model showed a significant positive relationship between the RCI of the lLM scores and the hemisphere that underwent surgery (p exact) and the age of onset of seizures (p exact) in a model that also accounted for sex, handedness, Engel scores, surgical technique and education (overall model R-squared and p). However, the lVM scores didn’t show a significant relation with other variables (Table 4).

**Table 4:**
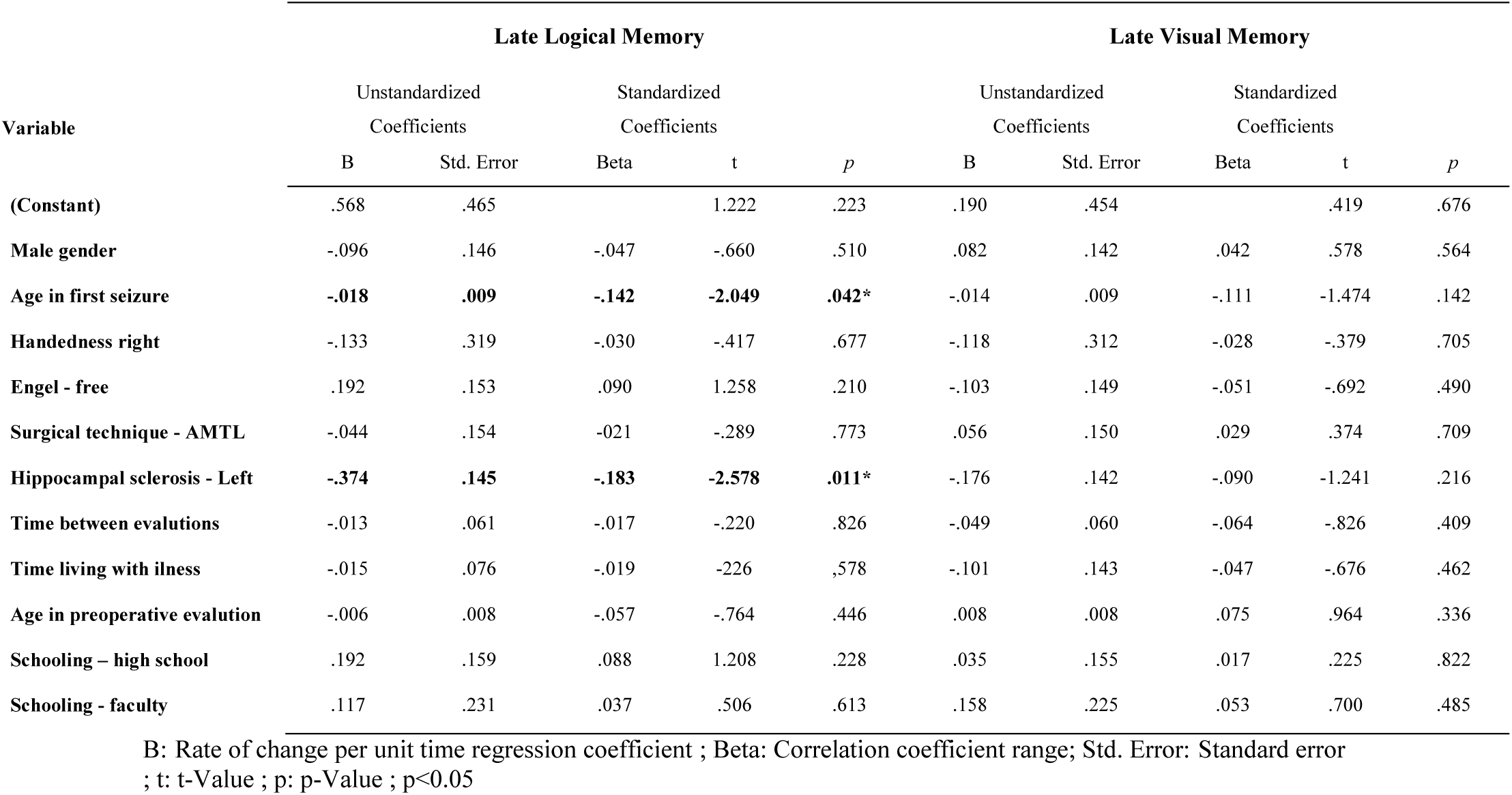
Multiple Linear Regression - Late Logical and Late Visual Memory.

**Late Logical-Verbal Memory:** the predictive factors for reliable change of late LVM are: operated hemisphere (p<0.05) and age of onset of seizures (p <0.05).

**Late Visual Memory:** no factor obtained statistical magnitude to be defined as a predictor for reliable change of late visual memory.

## DISCUSSION

When we divide our sample according to the lateralization of the disease and do not find statistically significant differences between the sociodemographic variables, we are ensuring a paired sample, with greater reliability in the identification of neuropsychological outcomes, since socio-educational factors, for example, could interfere with baseline cognitive and memory course outcomes, as previously described^24^. This division was made due to the large number of studies and scientific consensus that each cerebral hemisphere is responsible for specific primary brain functions, especially for distinct memory functions^25,26^.

In our sample, patients with epileptogenic focus in the LH presented preoperative and postoperative scores of immediate and late LM significantly lower when compared to the group of patients with focus in the Right Hemisphere (RH). This finding reinforces the knowledge that most right-handed individuals have cerebral hemispheric dominance for LM and language functions in the LH^27^. Our sample was substantially constituted by right-handed individuals, which would explain this phenomenon. On the other hand, patients with the disease in the RH did not present deficient baseline results for VM, which shows either a lower hemispheric dominance for this type of function or the importance of non mesial-temporal structures, coinciding with the results of a known previous study^27^. This way, it is necessary further investigations that delve into the knowledge regarding the specific cerebral location of the VM functions.

Patients who underwent resection in the RH obtained a statistically significant increase in late LM scores, typically associated with the left hemisphere^28^, after completion of surgery. This indicates an improvement in the contralateral cognitive function contralateral to that of the resected hippocampus. Still using this metric, patients who operated the LH did not present a statistically significant improvement on any of the two types of memory evaluated with this study. Alpherts et al^26^, in a long-term investigation of memory course in patients with MTL epilepsy, found similar results: individuals who operated the disease in the RH showed an improvement in memory in short time, but these results did not remain for a 6 years follow-up. However, the decline found in his patients who underwent resection in the LH lobe has progressed only in the first two years after resection and stabilized afterwards; later, the memory was stabilized. Another study confirmed this result, in which the memory decline occurred only in the first two years after surgery and in a reevaluation after 10 years there was a cognitive stability^29^. As our mean time of evaluation between the periods pre and postoperative was 2 and a half years, following the reasoning of the studies^26,29^, we can assume that the course of memory in relation to the exposed declines could also be attenuated. Nevertheless, this hypothesis could only be reliably confirmed from a reassessment of memory in a further follow-up.

Approximately 65% of our patients remained free of seizures at the time after the reevaluation. Our findings, coincide with the results of Helmstaedter^8^, in which 63% of the studied individuals had Engel I scores after surgery to treat MTL epilepsy. However, unlike the findings of this author, our study showed that being free of seizures is not a predictor of reliable improvement in memory; other investigations point to the same direction as our results^26,29^. Alvim et al^30^ demonstrated, in a structural neuroimaging study, that even after neurosurgery with removal of the epileptogenic focus, atrophy and progressive loss of gray matter exist, indicating that a mechanism underlying the pathology could be responsible for the decrease of brain volume even in patients free of seizures and in absence of cognitive decline. Investigations suggest there is a permanency of an epileptic spectrum even after the seizures are extinguished^30,31^. Pathology of hyperphosphorylated tau in the form of neuropilic wires, tangles and neurofibrillary pre-entanglements was found in a high proportion of individuals free of seizures following surgery, similar to those present in Alzheimer’s disease and Chronic Traumatic Encephalopathy (Braak III-IV staging)^32^.

The Reliable Change Index (RCI) method is understood by psychometrics as the most reliable to measure real changes after an intervention^18^. There was a larger percentage of reliable improvements for both for late LM and VM of patients of the rHS group, with a smaller decline in late VM scores. These results are also present in the study of Shah^33^, who used the same method in Indian patients with similar mesial temporal pathology as our sample: the individuals presented significant improvement of LVM when operated in the RH. Gül^34^ demonstrated in his research that the group of patients who operated the RH also showed a statistically significant improvement of VM, but these findings were not replicated in our investigation. However, according to the RCI, the proportion of patients who underwent neurosurgery in the RH and obtained a reliable improvement in late LVM and VM was larger in relation to the number of worsening. This phenomenon occurred inversely in patients who operated the LH: the percentage of worsening in both VLM and VM was larger than the number of improvements.

Multiple linear regressions showed that the LH and the age of onset of the first seizure were important predictors for reliable changes in LM after surgery. Patients that underwent a LH surgery had a lower rate of reliable change, that is, a tendency to decline in memory, which is already widely described in international literature specialized on the subject and can now be reproduced for the first time with the Brazilian population. Two studies^4,35^ performed with patients who underwent surgery to treat MTLE demonstrated that age at the procedure is a predictor of cognitive change: the younger the individual, the better the prognosis to the course of memory. In our research, these results were not evidenced, not even the time in which the individual lived with the disease before the neurosurgical treatment. However, the age of onset of the first seizure was a predictive factor: this occurrence could be explained by the theory of neural plasticity and cerebral reorganization^36^, since a younger brain would have a greater potential for reorganization of cognitive functions when exposed to a situation of disorganization for the first time. Thus, the lower the age of onset of seizures, the higher the score towards reliable memory improvement. No predictors were found for reliable visual memory change.

Because it is a retrospective cohort study, we found some limitations, such as the loss of participants due to inadequate filling of some charts of neuropsychological tests protocols. Also, a reevaluation of memory in a space with short time, average of 2 ½ years. As future perspective, we suggest follow-up of patients operated for a longer period, as well as the use of RCI to verify other cognitive functions in patients with mesial temporal lobe epilepsy and other types of epilepsy.

## CONCLUSION

After this study we can evidence the LH dominance for the functions of late LM in patients with refractory MTLE, with HS as the underlying disease. Also, a statistically significant improvement of both iLM and lLM, in a group way, in patients who underwent resection surgery of the epileptogenic focus on the RH. Patients with a RH lesion had a higher percentage of reliable improvement in both VM and LM scores. Having the epileptogenic focus in the LH and late onset age of the first seizures were shown to be predictive factors for a reliable worsening of LM. Thus, surgical treatment can be understood as an extremely effective alternative in MTLE patients with HS, since most of them become free of disabling seizures after the procedure and the smallest portion undergoes reliable changes of memory. The prognosis is even more positive when the disease occurs in the right cerebral hemisphere, where the functions of logical-verbal memory tend to be better in the first years after surgery.

## ETHICAL CONSIDERATIONS

This research complies with all the norms established by Law 466/2012 regarding Human Studies, according to the opinions issued by the Research Ethics Committees from UFRGS and PUCRS, respectively under the numbers: 2,471,665 and 2,492,372. The authors also state that this research was conducted in accordance with the principles of the World Medical Association Declaration of Helsinki.

## ACKNOWLEDGMENTS

The authors thank all the professionals who work in the Epilepsy Surgery Program of the São Lucas Hospital of PUCRS and the statistical advice of the same institution. This study was financed in part by the Coordenação de Aperfeiçoamento de Pessoal de Nível Superior – Brazil - CAPES (Coordination of Improvement of Higher Education Personnel) – Finance Code 001

